# Behavioural responses to video and live presentations of females reveal a dissociation between performance and motivational aspects of birdsong

**DOI:** 10.1101/624015

**Authors:** Logan S. James, R. Fan, J.T. Sakata

## Abstract

Understanding the regulation of social behavioural expression requires insight into motivational and performance aspects of social behaviours. While a number of studies have independently investigated motivational or performance aspects of social behaviours, few have examined how these aspects relate to each other. By comparing behavioural variation in response to live or video presentations of conspecific females, we analysed how variation in the motivation to produce courtship song covaries with variation in performance aspects of courtship song in male zebra finches (*Taeniopygia guttata*). Consistent with previous reports, we observed that male zebra finches were less motivated to produce courtship songs to videos of females than to live presentations of females. However, we found that acoustic features that reflect song performance were indistinguishable between songs produced to videos of females and songs produced to live presentations of females. For example, songs directed at video presentations of females were just as fast and stereotyped as songs directed at live females. These experimental manipulations and correlational analyses reveal a dissociation between motivational and performance aspects of birdsong and suggest a refinement of neural models of song production and control. In addition, they support the efficacy of videos to study both motivational and performance aspects of social behaviours.

## INTRODUCTION

The extent and quality of various social displays, including communicative and courtship behaviours, reflect an individual’s motivation and performance. Motivation refers to the “drive” to display a behaviour whereas performance refers to the fine motoric aspects of the behaviour. For example, internal and external states can affect the likelihood of displaying maternal behaviours (e.g., pup retrieval and grooming), and the latency and efficiency of pup-directed behaviours can vary between individuals as well as within individuals over time (Champagne et al., 2003; Clark et al., 2002; Stolzenberg et al., 2012). Both the motivation to engage in maternal behaviours and the performance of various components of maternal behaviour have important developmental consequences, and such findings highlight the importance of investigating both motivation and performance to gain a comprehensive understanding of social behaviour (Meaney, 2001; Rilling and Young, 2014). However, motivation and performance are often studied independently, and relatively little is known about the relationship between mechanisms regulating motivational and performance aspects of behaviour. In particular, little is known about the extent to which factors that affect the motivation to display a behaviour similarly affect the performance of the behaviour.

Birdsong provides an excellent opportunity to assess the degree to which mechanisms underlying motivational and performance aspects of social behaviour are shared or independent. When presented with an adult female, adult male songbirds become motivated to produce courtship song, and individual differences in this motivation is important because female songbirds also tend to prefer males that display greater motivation to sing (Bradbury and Vehrencamp, 2011; Catchpole and Slater, 2008; Gil and Gahr, 2002; Sakata and Vehrencamp, 2012). Further, males alter a number of song performance features when producing courtship songs compared to non-courtship songs (Chen et al., 2016; Moser-Purdy and Mennill, 2016; Sakata and Vehrencamp, 2012; Toccalino et al., 2016; Vignal et al., 2004; Woolley and Kao, 2015). For example, male zebra finches produce songs that are faster and more acoustically stereotyped when courting female conspecifics than when singing in isolation (Chen et al., 2016; Cooper and Goller, 2006; Kao and Brainard, 2006; Sossinka and Böhner, 1980; Woolley et al., 2014). These performance-related song traits can affect a male’s attractiveness and reproductive success, since female songbirds prefer the courtship version of an individual male’s song, as well as males with song features that are generally characteristic of courtship song (e.g., faster and longer songs; Gil and Gahr, 2002; Podos et al., 2009; Woolley and Doupe, 2008).

Despite knowledge about the functional relevance of motivational and performance aspects of birdsong, little is known about how experimental variation in the motivation to produce courtship song relates to experimental variation in song performance. Brain areas that underlie the motivation to sing project to sensorimotor brain regions that regulate song performance, suggesting that song motivation could influence song performance (reviewed in Riters, 2012; Riters et al., 2004; Woolley and Kao, 2015). In addition, seasonal changes in the motivation to produce song have been found to covary with seasonal changes in song performance (Smith et al., 1997; Smith et al., 1995). On the other hand, some studies have found a dissociation between song motivation and performance (Alward et al., 2013; Ritschard et al., 2011; Toccalino et al., 2016).

Here, we investigated variation in vocal performance across conditions that are known to modulate the motivation to produce courtship song. Video playbacks of social stimuli have been used to elicit a wide range social behaviours (Evans and Marler, 1991; Fleishman and Endler, 2000; Gonçalves et al., 2000; Guillette and Healy, 2017; Oliveira et al., 1999; Ophir et al., 2005; Ord et al., 2002; Rosenthal, 1999; Uetz and Roberts, 2002; Ware et al., 2016), including courtship song in songbirds (Galoch and Bischof, 2007; Ikebuchi and Okanoya, 1999; Takahasi et al., 2005). Despite that videos can elicit courtship song, male songbirds have been found to sing less to videos of females than to live presentations of females (Ikebuchi and Okanoya, 1999). However, it is not known whether performance aspects of courtship song (e.g., tempo and stereotypy) are similarly reduced for songs produced to video presentations of females. Previous studies of other social behaviours have found that behavioural performance can be distinct when individuals are presented with video or live presentations of conspecifics (Balshine-Earn and Lotem, 1998; Ord et al., 2002; Swaddle et al., 2006). Consequently, we analysed motivational and performance aspects of male zebra finch song in response to video and live presentations of females.

## MATERIALS AND METHODS

### Animals

Adult male zebra finches (*Taeniopygia guttata*; >4 months; n=13) were bred and raised in our colony at McGill University. Males were socially housed in same-sex group cages and visually isolated from females. Birds were kept on a 14L:10D photoperiod, with food and water provided *ad libitum*. All procedures were in accordance with McGill University Animal Care and Use Committee protocols, as well as guidelines from the Canadian Council on Animal Care.

### Video stimuli

Stimulus females were videotaped using a SONY DCR-SR 220 HD camcorder at 60 frames per second. We gathered footage of individual females perched at camera-level in front of a neutral background. Adobe Premiere 2017 was used for minor white balance corrections, cropping and trimming. Playback clips were 30 seconds in duration and featured a silent, perched female engaged in a moderate level of activity (e.g., movements of head and along perch but no flying; Movie 1). A total of six females were filmed with three clips created per individual.

### Behaviour testing and song collection

Figure 1 illustrates the experimental setup used during song collection. Male finches were isolated in individual cages (20 × 20 × 20 cm) inside sound-attenuating chambers (“soundboxes”; TRA Acoustics, Ontario, Canada) from at least one day prior to experiments. All songs were recorded using an omnidirectional microphone (Countryman Associates, Inc., Menlo Park, CA, USA) positioned directly above the male’s cage. During experiments, song was detected and digitized using a sound-activated system (Sound Analysis Pro v.1.04 http://ofer.sci.ccny.cuny.edu/html/sound_analysis.html, digitized at 44.1 kHz). A Microsoft Surface Pro 3 tablet (2160 × 1440 pixels) was used to playback videos and was fixed to a wall of the soundbox. The tablet was placed in the soundbox at least 10 minutes before the onset of testing and was positioned ∼12 cm from the male’s cage. We sized the video playback window such that stimulus bird in the video was approximately life-size at the distance between the cage and tablet. The screen was blank (black) when not displaying video stimuli. A camera mounted above the tablet provided a live stream of the experimental bird for monitoring. All experiments began within 2 hours of lights turning on.

**Figure 1:**
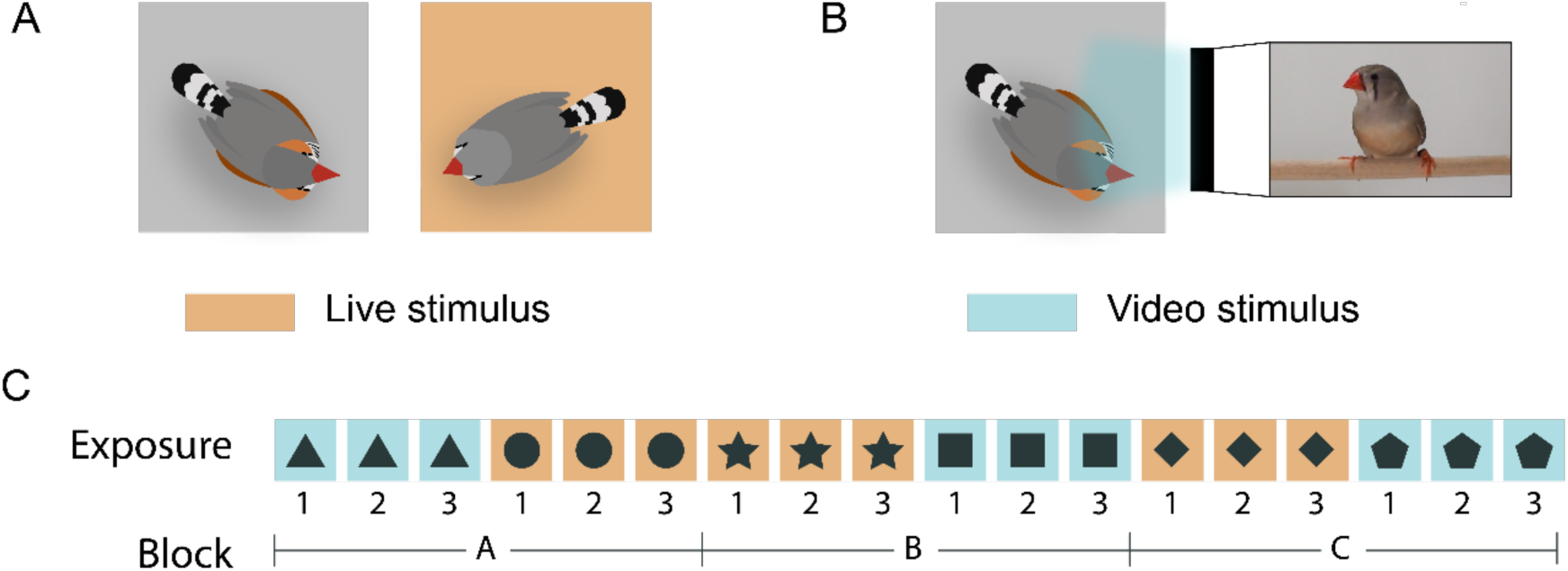
Experimental designs for live and video presentations of females. Schematics represent experimental setup during (A) exposure to live female stimulus and (B) exposure to videos of females (not drawn to scale). (C) Example of stimulus presentation order for one male. Males were tested in three blocks (A, B, C), with each block consisting of three consecutive exposures to distinct videos of an individual female and three consecutive live exposures to an individual female. All presentations were separated by a five-minute interval. Shapes represent stimulus individuals.

During experiments, we collected courtship songs from male zebra finches using a design similar to that described by Toccalino et al. (2016). Specifically, each male was briefly exposed to six different females, three via live presentations and three via video presentations. During live female presentations, an experimenter opened the soundbox door and placed a cage housing a conspecific female next to the experimental male’s cage. The soundbox door was then closed, and the female remained in the soundbox for 30 seconds. During video presentations of females, an experimenter opened the soundbox door, started a video of a female, and then closed the door to the soundbox. Videos played for 30 seconds and ended on a black screen.

Males were exposed to a total of 6 randomly-chosen stimulus females from a pool of 6 videotaped females and 12 live females. Females that were videotaped were distinct from those used for live presentations. Video and live presentations were grouped into three blocks (blocks A-C; Figure 1C), with each block consisting of three consecutive exposures to either a video or live presentation of an individual female (exposures 1-3), followed by three consecutive exposures to the other stimulus type. Within each block of video presentations, males were exposed to distinct video clips of the same female. All presentations were separated by five-minute intervals. The order of conditions (video vs. live presentation) within a block was pseudo-randomly determined to balance the order of conditions. The first condition presented in each block was determined by a coin flip, and, if the first conditions of the first two blocks were the same (e.g., video first for blocks A and B), the order was reversed for the last block, ensuring that no experimental session consisted of blocks that each started with the same condition.

We categorized a male’s song as directed toward the live or video presentation of a female if at least two of the following conditions were met during song production: (1) the male approached or oriented toward the stimulus females; (2) the male fluffed his plumage; and (3) the male pivoted his body from side to side (James and Sakata, 2015; Kao and Brainard, 2006; Morris, 1954; Toccalino et al., 2016). Typically, male zebra finches produce courtship song within a few seconds of stimulus presentation. Males in this study each produced a minimum of three courtship songs to video or live presentations of females (18.0 ± 3.3 and 10.0 ± 2.1 song bouts per male, respectively, toward live and video presentations of females).

We also collected non-courtship, or undirected (UD) songs (i.e., songs produced spontaneously when alone) during the experiment to contrast with courtship songs. Undirected songs were generally produced during the five-minute intervals between female exposures. In cases where few UD songs were produced between female presentations, UD songs produced in the 30 minutes before and after the testing period were used for analysis (16.2 ± 2.9 UD song bouts per male)(e.g. James and Sakata, 2015; Sakata et al., 2008; Toccalino et al., 2016).

### Song analysis

We used the following definitions for our analyses (Figure 2). “Song bouts” are defined as epochs of singing that are separated by at least 1 s of silence (e.g. Johnson et al., 2002; Poopatanapong et al., 2006). Each song bout consists of a stereotyped sequence of vocalizations called a “motif” that is repeated throughout the bout (Sossinka and Böhner, 1980; Zann, 1996). Motifs consist of distinct vocal elements (“syllables”) that are separated by at least 5 ms of silence. The first motif of a bout is preceded by repetitions of brief vocal elements called “introductory notes.”

**Figure 2:**
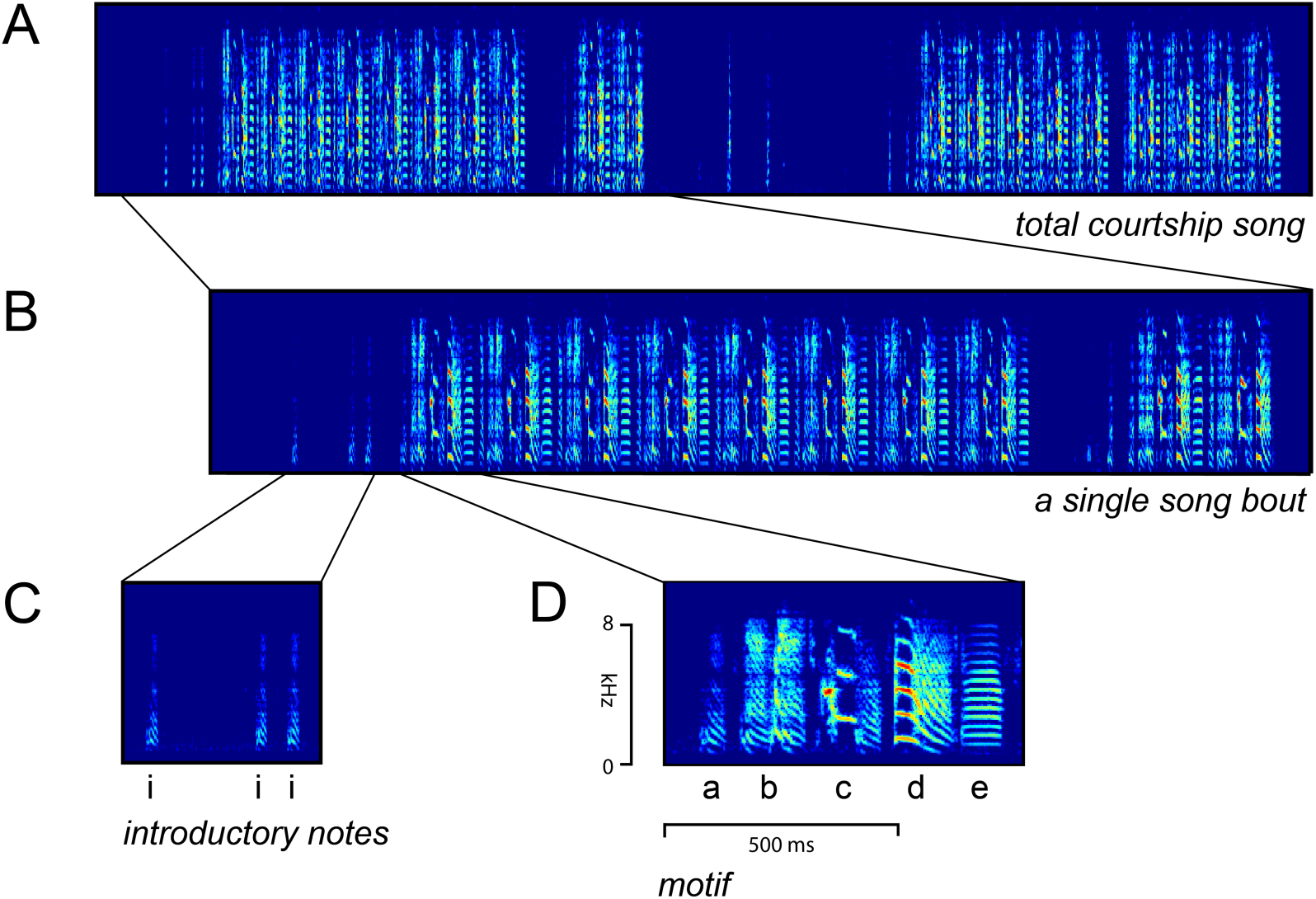
Organization of zebra finch song. Spectrograms plot the frequency (kHz) against time (ms) with brightness reflecting amplitude. (A) An example of song produced during a 30-sec exposure to a video. In this example, the male produced two song bouts (bouts separated by ≥ 1 second of silence). (B) Zoomed-in image of a single song bout; song bouts begin with the repetition of introductory notes (C) and consist of a stereotyped sequence of syllables (‘motif’) that is repeated throughout the bout (D). The motif of this bird consists of five separate syllables, each separated by ≥ 5 ms of silence.

Our primary measure of courtship song motivation was the total amount of time (seconds) males engaged in courtship song across all exposures to live or video presentations of females (“time spent singing”). We also deconstructed this measure into various components, including the likelihood that males will produce courtship song on a given exposure and the total duration of song during each exposure. We also broke down the total song duration during each exposure into the number of bouts produced during each exposure, and the duration of each of those bouts. Bout durations were defined as the interval between the onset of the first syllable to the onset of the last syllable of the bout.

We analysed song features that are consistently affected by social stimuli and that have been used as indices of song performance (Sakata and Vehrencamp, 2012). In particular, we measured the number of introductory notes preceding song, song tempo, and the variability of the fundamental frequency (FF) of syllables with flat, harmonic structure (Chen et al., 2016; Cooper and Goller, 2006; James and Sakata, 2014; Kao and Brainard, 2006; Sakata et al., 2008; Stepanek and Doupe, 2010). For these analyses, we first manually labelled syllables and introductory notes following amplitude-based element segmentation using custom software written in MATLAB (The MathWorks, Natick, MA, USA). Introductory notes were quantified by starting with the note immediately preceding the first syllable of the bout and counting backwards until we reached ≥1 second of silence. Motif duration was defined as the duration from the onset of the first syllable of the motif to the onset to the last syllable of the motif and was used as the metric for song tempo (e.g., James and Sakata, 2015; Kao and Brainard, 2006; Sakata et al., 2008). We restricted the analysis of song tempo to the first motif of the bout, because motif durations have been found to change across the song bout and because bout durations differ between live and video presentations (see Results; Chi and Margoliash, 2001; Cooper and Goller, 2006; Glaze and Troyer, 2006). Finally, we computed the fundamental frequency (FF) of syllables with flat, harmonic structure (e.g., syllables ‘c,’ ‘d,’ and ‘e’ in Figure 2D) by calculating the autocorrelation of a segment of the sound waveform and measuring the distance (in Hz) from the zero-offset peak to the highest peak in the autocorrelation function. We measured the FF for every rendition of the syllable, and then computed the coefficient of variation (CV: standard deviation/mean) per syllable per condition as an index of acoustic variability (Sakata et al., 2008; Toccalino et al., 2016). We also computed these same measures of song performance for UD songs to contrast with performance for songs directed at videos of females [video-directed (VD) song] and songs directed at live females [live-directed (LD) songs].

### Data Analysis

We compared song motivation between experimental conditions for all 13 males. However, five males produced courtship songs only during live presentations of females; therefore, in our direct comparisons of song motivation and performance during live and video presentations of females, data were restricted to the eight males that produced songs during both live and video presentations of females. Data for a number of song features were computed for each exposure (total song duration per exposure, bout duration, first motif duration, introductory notes and fundamental frequency), and for these data, we only analysed exposures during which song was produced.

Statistical analyses were conducted in R 2.15.1. We used linear mixed models (LMMs) and generalized linear mixed models (GLMMs) within the ‘lme4’ library (Bates et al., 2015) to compare singing behaviour across experimental conditions. Our experimental design consisted of three testing blocks (blocks A-C), with each block consisting of three consecutive exposures to videos of a single female and three consecutive exposures to a live female (exposures 1-3; Figure 1C). Therefore, we ran three-way factorial models with Block (A-C; ordinal), Exposure (1-3; ordinal), Condition (live vs. video; nominal) and all possible interactions as fixed effects. Because of the repeated-measures nature of this design, we also included Bird ID as a random factor. Further, because birds can produce multiple bouts within an exposure and because bout number can affect some song features (see Results), we also ran four-way full-factorial models with the same three fixed effects plus Bout (1-3; ordinal).

In the analysis of the likelihood to produce courtship song, we had one binary response variable (whether the bird produced at least one courtship song bout or not during each exposure); therefore, we ran this model as a GLMM with a binomial error family. The number of song bouts produced during each exposure and the number of introductory notes preceding song bouts (see above) were count responses; consequently, we ran these models as GLMMs with a Poisson error family. Total time spent singing, total song duration per exposure, and song bout durations were highly skewed; therefore these data were analysed with a gamma error family and a log link (data plotted following log-transformation for ease of presentation). Finally, the total number of exposures in which at least one song bout was produced and the duration of the first motif were analysed with LMMs with a Gaussian error family. Prior to running the statistical models, data were visually screened to assess model fit using Q-Q plots. To test the significance of each mixed model, we ran Type II Wald chi-square tests using the ‘car’ library (Fox et al., 2011).

We used a different statistical model to analyse experimental variation in the CV of FF. This is because the CV (standard deviation divided by the mean) cannot be computed for a single rendition and needs to be computed across multiple renditions of syllable (or motif). To provide reliable estimates of the CV of the FF of a particular syllable, we measured the CV across each rendition of a syllable for all songs produced during all blocks and exposures. The statistical model to analyse variation in the CV of FF also differs from the ones described above because birds can produce multiple syllables for which we calculated the FF (n=13 syllables across the eight males). Consequently, for the analysis of the CV of FF, we ran a LMM with a Gaussian error family, and with Condition as the fixed effect and Syllable ID nested in Bird ID as a random effect so that we could directly compare the CV of the same syllable across conditions.

In addition to assessing differences in song performance across VD and LD songs, we also compared song performance of VD and LD songs with those of UD songs. For these analyses, we computed data for VD and LD song across all renditions of song (i.e., across blocks, exposures, and bouts) and compared these values to those for UD song. We ran one-way models with similar parameterization as before, with Condition as the sole independent variable. We used a Poisson error family for introductory notes and a Gaussian error family for first motif durations and CV of FF. Bird ID was a random variable for introductory notes and first motif durations, and Syllable ID nested in Bird ID was a random effect for CV of FF. For these analyses, we ran Tukey’s tests with the Holm correction using the ‘multcomp’ library (Hothorn et al., 2008) for post-hoc contrasts across the three conditions. As with the previous analysis, we nested Syllable ID within Bird ID as the random effect in the analysis of the CV of FF.

To gain further insight into the relationship between song changes across experimental conditions, we also analysed the extent to which motivational and performance changes driven by video presentations of females co-varied with motivational and performance changes driven by live presentations of females. We correlated the total amount of song a male produced to live presentations of females with the total amount of song a male produced to video presentations of females. In addition, we computed the percent change of song features from UD to VD song and from UD to LD song, and correlated these percent changes. Because we measured the CV of FF of multiple syllables, and each syllable within a bird could change independently, we ran these correlations as LMMs with the Gaussian error family with Bird ID as a random effect. All other relationship were analysed using Pearson’s product-moment correlations.

Finally, we analysed the extent to which individual variation in the differential motivation to produce courtship song to video and live presentations of females covaried with individual variation in the differential modulation of song performance across video and live presentations. Specifically, we correlated the difference in time spent singing LD and VD song with the difference in the modulation of each song feature from UD to LD and UD to VD. We used Pearson’s correlations for analyses of the number of introductory notes and the first motif duration. Because the differential motivation to produce courtship song to videos or live presentations of females is summarised by one value per bird, we calculated the average change in the CV of FF across all syllables produced by each bird to relate performance to motivation (i.e., each bird has only one data point representing the average percent change in the CV of FF of syllables).

## RESULTS

### Differences in the motivation to produce courtship song to video vs. live presentations of females

We first counted, for each male (n=13), the number of exposures to live females or videos of females in which a male produced at least one courtship song (out of nine exposures per male for each condition). We found that male zebra finches produced courtship song on significantly more exposures to live females (6.6 ± 0.7 (out of 9); mean ± SEM) than to videos of females (3.6 ± 0.9 (out of 9); χ ^2^_1_=19.0, p<0.0001). Upon further inspection, we noted that, while most birds produced courtship songs to both live and video presentations of females (n=8), five birds sang exclusively towards live females (no males sang exclusively towards videos of females). However, the number of exposures with courtship song remained significantly higher for live presentations of females even when analyses were restricted to males that produced courtship song to both video and live presentations of females (live: 7.8 ± 0.8; video: 5.9 ± 0.8; χ ^2^_1_=8.6, p=0.0034).

The total amount of time a male spends singing towards a female is widely considered a reliable measure of song motivation. As such, we compared the total amount of time male zebra finches sang to live and video presentations of females (i.e., total duration of song across all exposures). Because this study is focused on experimental differences in motivation and performance features and because performance features of video-directed songs cannot be computed for males that did not sing to videos of females, we limited our analyses to birds that sang in both conditions (n=8). Overall, we found that birds produced significantly more song towards live females (116 ± 18 seconds) than to videos of females (46 ± 8 seconds; χ ^2^_1_, p<0.0001; Figure 3A; Table S1).

**Figure 3:**
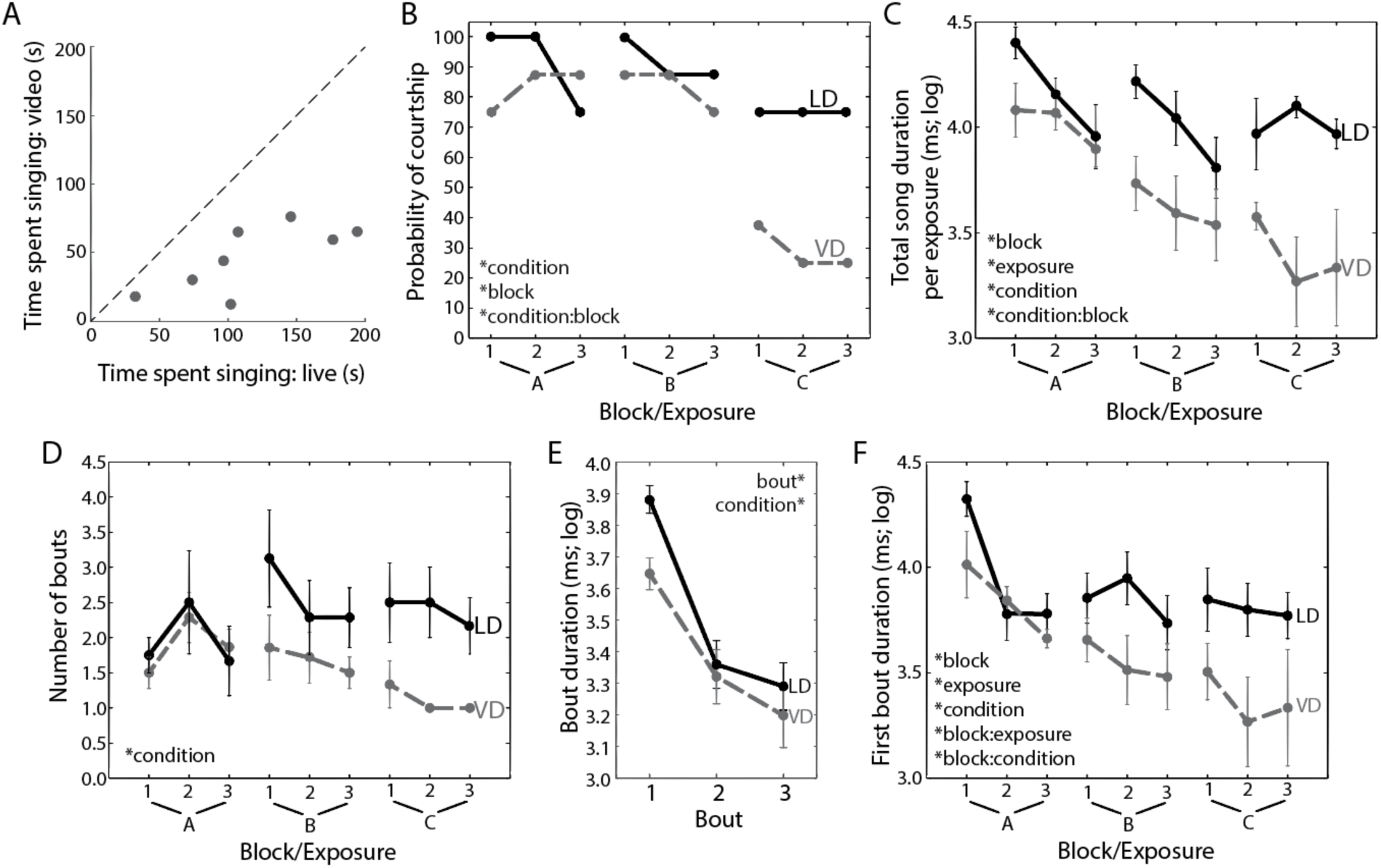
Male zebra finches produced courtship song to both video and live presentations of females but were less motivated to produce song to video presentations of females. For panels (B-F), data for video-directed (VD) and live-directed (LD) songs are plotted, respectively, with dashed grey and solid black lines. All error bars are the standard error of the mean. (A) Birds produced more song towards live females than towards videos of females. Each dot represents an individual bird, and the dashed line depicts the line of unity. (B) The probability of courtship per exposure is affected by condition, block and a marginally significant interaction between condition and block. (C) Total song duration per exposure is affected by condition, block, exposure and the interaction between condition and block. (D) The number of song bouts produced during each exposure to a female is only affected by condition. (E-F) Song bout duration is affected by bout, block, exposure, condition and the interaction between block, exposure and bout (see Results). For ease of presentation, we first depict the effects of bout and condition (E), followed by the effects of block, exposure and condition on the duration of the first bout produced during each exposure (F).

Differences in the total amount of song produced to live vs. video presentations of females could be caused by a number of factors, including variation in the probability of producing courtship song on each exposure, in the total amount of song on each exposure, in the number of song bouts produced on each exposure, and in bout durations. We first analysed whether males differed in their probability of producing courtship song towards live or video presentations of females on each individual exposure to a female stimulus. We performed a 3-way GLMM with Condition (live or video), Block (A-C) and Exposure (1-3; ordinal) as independent factors, Courtship (0 or 1; binomial) as the response variable, and Bird ID as a random effect (all birds included in this analysis). We found significant effects of Condition (χ ^2^_1_, p=0.0323) and Block (χ ^2^_2_, p=0.0122), indicating that birds were significantly more likely to produce courtship song to live females than to videos of females and that the likelihood of a male producing courtship song decreased across blocks (Figure 3B). In addition, there was a marginally significant interaction between Condition and Block (χ ^2^_2_ p=0.0588), with differences across conditions being larger for later blocks.

To further reveal the factors that contributed to the overall difference in the amount of courtship song produced to live vs. video presentations of females, we examined the total amount of courtship song produced (in seconds) during each video or live exposure to a female (each exposure to a female stimulus was 30 sec in duration; Table S1). We found significant effects of Condition(χ ^2^_1_, p<0.0001), Block (χ ^2^_2_, p<0.0001), and Exposure (χ ^2^_2_,p<0.0001) on total song duration per exposure (Figure 3C). Overall, song durations per exposure were shorter in response to video presentations of females than live presentations, and durations decreased across blocks and exposures. In addition, there was a significant interaction between Condition and Block (χ ^2^_2_, p<0.0001), which was characterized by smaller changes in song durations across blocks for live presentations than for video presentations. As such, the difference in song durations between video and live exposures became larger over the blocks of testing.

Because birds produce courtship songs in bouts (i.e., epochs of song separated by ≥1 sec of silence), differences in courtship song duration per exposure could be due to differences in the number of song bouts produced during each exposure as well as differences in the lengths of song bouts. Consequently, we first analysed the number of bouts that male zebra finches produced on each exposure (Supplementary Table S1). We found a significant effect of Condition (χ ^2^_1_=5.2, p=0.0221; Figure 2D), with males producing fewer song bouts per exposure to videos of females (mean ± SEM: 1.70 ± 0.12 bouts) than per exposure to a live female (2.32 ± 0.18 bouts). While the interaction between Condition and Block was not statistically significant, visual inspections of the data indicate a trend for the difference between video and live presentations to become larger over the blocks of testing.

To analyse song bout duration, we ran a four-way factorial mixed effects model with the same fixed factors as above (Condition, Block and Exposure) as well as Bout (i.e., the serial order of bouts within each exposure; ordinal). Because birds rarely produced more than three bouts in an exposure, we limited our analysis to the first three bouts per exposure (models were rank deficiency when data for all bouts were included; Table S1). We found significant effects for all main factors (Condition: (χ ^2^_1_=40.3, p<0.0001; Block: (χ ^2^_2_, =46.5, p<0.0001; Exposure: (χ ^2^_2_=21.0, p=<0.0001; Bout: χ ^2^_1_=129.6, p<0.0001), as well as a three-way interaction between Block, Exposure, and Bout (χ ^2^_8_=36.5, p<0.0001) and between Exposure, Condition and Bout (χ ^2^_4_=9.8, p=0.0431; Figure 3E). We also observed a significant interaction between Block and Exposure (χ ^2^_4_=10.6, p=0.0318) and a marginal interaction between Block and Bout (χ ^2^_4_=8.4, p=0.0795). Overall, bout durations were longer for songs produced to live presentations of females than for songs produce to video presentations of females. Additionally, bout durations decreased across blocks, across exposures within blocks, and across bouts produced within each exposure to a female stimulus.

Because of the complexity of the four-way model, we conducted another analysis limited to the data from the first bout (i.e., Bout not included as an effect in the model) to obtain a simplified depiction of variation in song bout duration (Table S1). We observed significant main effects for all three factors (Condition: χ ^2^_1_=38.7, p<0.0001; Block: χ ^2^_2_=34.1, p<0.0001; Exposure: χ ^2^_2_=38.7, p=0.0001) as well as significant interactions between Block and Exposure (χ ^2^_4_=18.3, p=0.0011) and between Block and Condition (χ ^2^_2_=9.9, p=0.0071; Figure 3F). Overall, the duration of the first bout of courtship song was longer for songs produced to live presentations of females than for songs produced to video female presentations, and bout durations became shorter across blocks and exposures. The interactions were characterized by larger decreases across exposures during block A than during blocks B and C, and by larger decreases across blocks for video presentations than for live presentations.

Together, these analyses indicate that differences in total amount of courtship song in response to video and live presentations of females were due to differences in the likelihood of producing courtship song, the number of song bouts per exposure, and the duration of individual song bouts.

Despite differences in the amount of courtship song produced to live vs. video presentations of females, it is possible that individual variation in the motivation to produce courtship songs to videos of females is related to variation in the motivation to produce courtship songs to live presentations of females. Therefore, we correlated individual variation in the total amount of time males spent singing to live and video presentations of females. Consistent with the notion that motivation to court videos of females is related to the motivation to court live females, we found a significant correlation between the total amount of song produced to live versus video stimuli (n=8; r=0.73, p=0.0382).

### Lack of differences in performance features of courtship songs produced to video vs. live presentations of females

Our results support that zebra finches are less motivated to court videos of females than live females, and we next sought to determine whether performance aspects also varied across songs directed at live or video presentations of females. To this end, we compared various measures of song performance (see Methods) between video-directed (VD) and live-directed (LD) songs among males that produced both types of songs (n=8 birds).

To analyse differences in the number of introductory notes preceding song, we first ran a four-way factorial model with Condition, Block, Exposure and Bout (limited to the first three bouts; see above) as fixed effects, Bird ID as a random factor, and the number of introductory notes before each bout of song as a Poisson response variable (Table S2). Importantly, we found no significant effect of Condition or interaction between Condition and other variables for the number of introductory notes. We only observed an effect of Bout (χ ^2^_6_=123.9, p<0.0001), with the number of introductory notes decreasing across consecutive bouts produced during an exposure to a stimulus (Figure 4A). We also ran a similar analysis with data limited to the first bout of song on each exposure (Bout excluded as a factor) and, again, found no significant variation across conditions, blocks, and exposures (Figure 4B).

**Figure 4:**
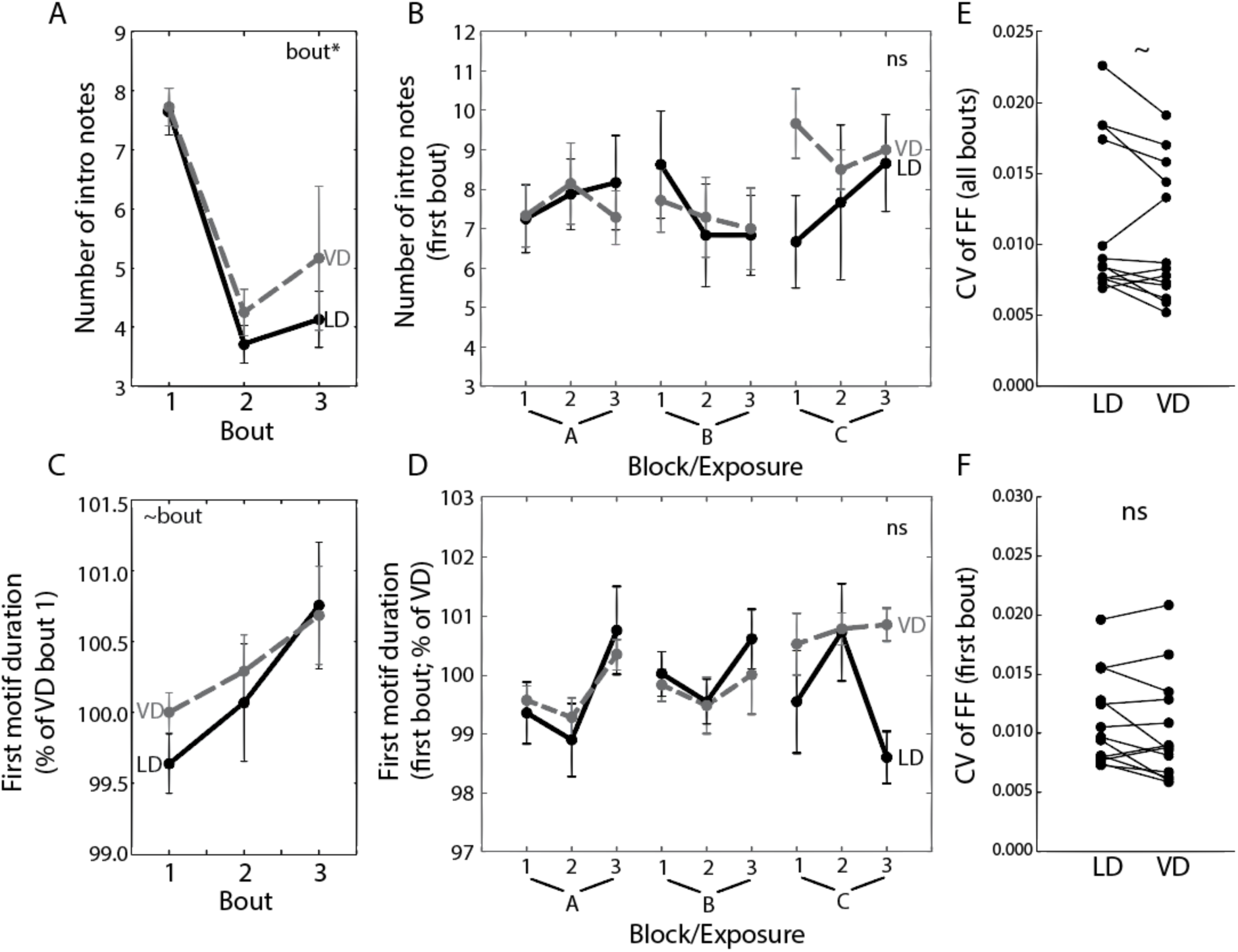
Male zebra finches produce video-directed (VD) and live-directed (LD) courtship songs with similar song features. For panels (A-D), data for VD and LD songs are plotted with dashed grey and solid black lines, respectively. For ease of presentation we first depict the effects of bout and condition for the first three bouts (A, C), followed by the effects of block, exposure and condition on a data set limited to the first bout produced (B, D). All error bars are the standard error of the mean. (A) The number of introductory notes is affected by bout, with no significant difference between VD and LD song. (B) The number of introductory notes before the first bout of song is not affected by block, exposure or condition. (C) There was a marginal effect of bout on the first motif duration, but no significant effect of condition. Data in the figure are normalized to the mean duration of VD song for data visualization purposes (see Methods). (D) The first motif duration of the first bouts of song produced during an exposure was not significantly affected by block, exposure or condition. (E, F) The CV of FF was marginally lower for VD song than LD song when all bouts were included in the analysis (E), but there was no significant difference when only the first bouts per exposure were analysed (F). For (E, F) each pair of connected dots depicts one syllable within a bird (n=13 syllables across the eight birds).

To analyse variation in song tempo between VD and LD song, we calculated the duration of the first motif of each song bout and analysed experimental variation in first motif durations using the same four-way factorial model as above (only first three bouts). Only the first motif in each bout was analysed for this comparison because motif durations change as bout length increases (e.g., Chi and Margoliash, 2001; Glaze and Troyer, 2006; James and Sakata, 2014; James and Sakata, 2015) and because bout lengths differed between VD and LD song (Figure 3; Table S2). There was no significant effect of any factor, including Condition, on song tempo (Figure 4C). We also ran a three-way factorial model using only data from the first bout produced per exposure and, again, found no significant effects (Figure 4D).

The FF of syllables with flat, harmonic structure is less variable from rendition-to-rendition when males direct song at females (Sakata and Vehrencamp, 2012; Woolley and Kao, 2015). We calculated the CV of FF across all syllable renditions in every bout of song and compared this variability between conditions (i.e., Condition is the only independent variable). We found a marginally significant difference between Conditionsχ ^2^_1_=3.7, p=0.0545; Figure 4E) with VD song tending to have lower CVs than LD song. However, no significant difference between VD and LD song was observed when only data from the first bout were analysed (χ ^2^_1_=2.3, p=0.1266; Figure 4F). This difference in the magnitude of differences between VD and LD song is primarily due to a decrease in the CV of FF for LD song.

### Courtship songs produced to video or live presentations of females are distinct in performance from undirected song

Overall, the preceding analyses indicate a lack of difference between VD and LD songs for three performance measures: the number of introductory notes, song tempo, and spectral variability. However, these analyses do not indicate whether males alter VD songs to be distinct from non-courtship songs (undirected or UD songs) in the same way that LD songs differ from UD songs. We therefore compared performance measures of every VD and LD song of a male to all his UD songs. We found a significant effect of Condition for introductory notes (χ ^2^_2_=51.3, p<0.0001) with post-hoc contrasts indicating that both VD and LD songs were preceded by more introductory notes than UD songs and that VD songs were preceded by more introductory notes than LD songs (p<0.003 for all). We also found a significant effect of Condition on first motif duration (χ ^2^_2_=39.7, p<0.0001), with post-hoc contrasts indicating that motifs were shorter in VD and LD songs relative to UD songs (p<0.0001 for both). Finally, we found a significant effect of Condition on the CV of FF (χ ^2^_2_=8.8, p=0.01201) with post-hoc contrasts indicating that the CV of FF was lower for VD songs compared to UD songs (p=0.0089).

The preceding analyses indicated that VD songs were distinct from UD songs, with mixed results regarding LD songs. However, our analyses above highlight how performance features can change across bouts and how the number of bouts produced per exposure differed between video and live presentations of females (Figure 4); consequently, the previous results are confounded by experimental variation in the number of bouts per exposure. To examine variation without this confound, we conducted the same analyses with data restricted to the first bout of song per exposure. In addition, we limited our UD song data to songs preceded by at least 30 s of silence to approximate the first bout restriction for VD and LD songs (see Methods). The number of introductory notes was significantly affected by Condition (χ ^2^_2_=67.4, p<0.0001; Figure 5A), and this difference was due to the fact that VD and LD songs were preceded by more introductory notes than UD song (p<0.0001 for each). The duration of the first motif was significantly different across Conditions (χ ^2^_2_=34.3, p<0.0001; Figure 5B) and was shorter during VD and LD songs than during UD song (p<0.0002 for each). Finally, the CV of FF was also affected by Condition (χ ^2^_2_=12.2, p=0.0023; Figure 5C), with the CV of FF being significantly lower during VD song than during UD song (p=0.0015) and with the same trend for LD song (p=0.0875). Consistent with the analysis described above, all these song features were not significantly different between VD and LD songs. Taken together, these data indicate that males change their song performance when directing songs at video presentations of females and that the nature and degree of these changes are comparable between VD and LD songs. These data also suggest that shared mechanisms could mediate the modulation of courtship song in response to video and live presentations of females.

**Figure 5:**
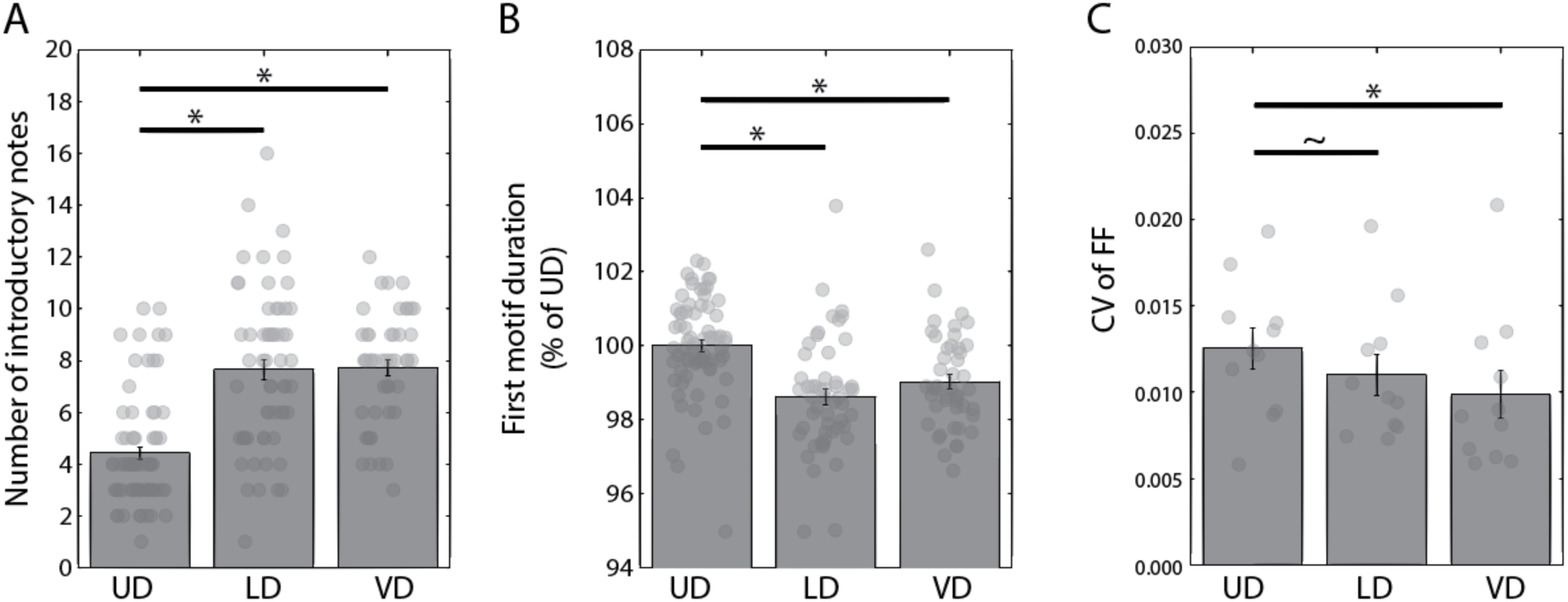
Temporal and spectral features of video-directed (VD) and live-directed (LD) songs were significantly different from those of undirected (UD) songs but not different from each other. (A) The number of introductory notes was significantly greater before VD and LD song than before UD song. (B) First motif durations were significantly shorter during both VD and LD song than during UD song. (C) The coefficient of variation (CV) of fundamental frequency (FF) was lower during VD and LD song than during UD song. Data were taken from the first bout of song for VD and LD song. “*” denotes p<0.05, and “∼” denotes p<0.10 (Tukey’s HSD test with Holm correction).

To further investigate whether the modulation of VD and LD songs could be mediated by similar neural mechanisms, we correlated individual variation in the magnitude of change in song performance from UD song to VD song with individual variation in the magnitude of change from UD song to LD song (data from first bout only). If shared mechanisms underlie the modulation of VD and LD song, we should observe significant and positive relationships between the magnitudes of song modulation for VD and LD song. The relationship was significantly positive for the number of introductory notes (r=0.72, p=0.0430; Figure 6A) and the CV of FF (χ ^2^_1_=14.4, p=0.0015; Figure 6C). The relationship was positive but not statistically significant for first motif duration (r=0.53, p=0.1766; Figure 6B).

**Figure 6:**
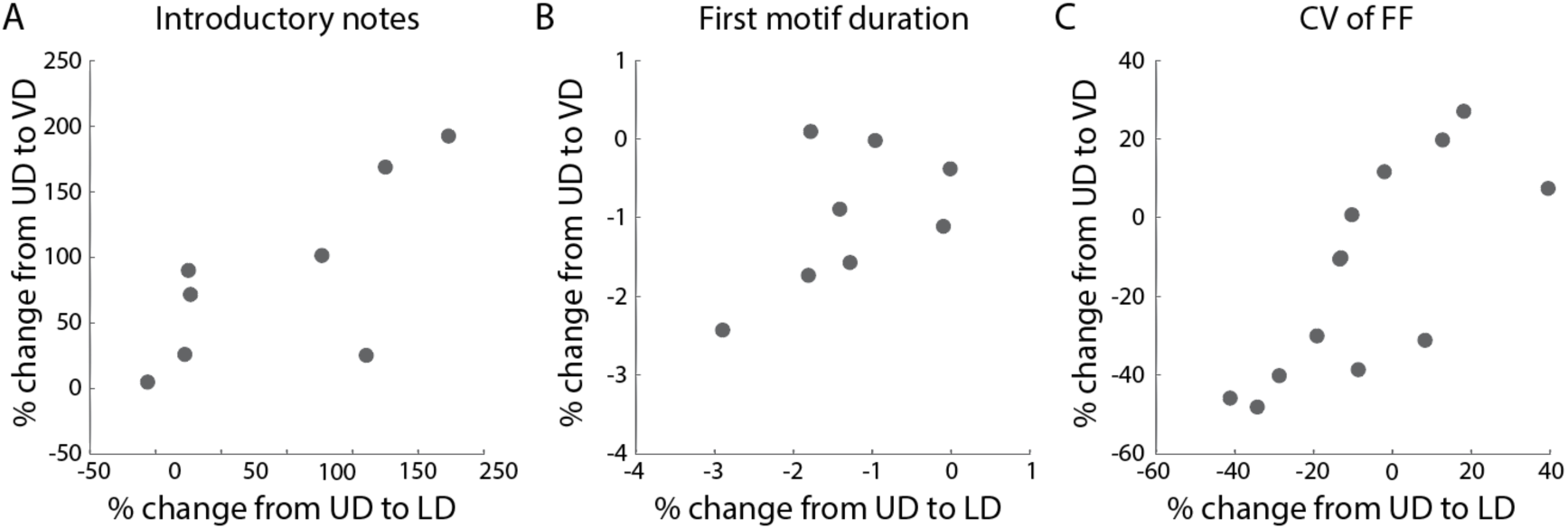
Relationships between the magnitudes of song changes from undirected (UD) to live-directed (LD) song (x-axis) and from UD to video-directed (VD) song (y-axis). Correlations were positive for all song features and were statistically significant for the number of introductory notes (A) and the CV of FF (C) but not for the duration of the first motif (B). For (C), each dot represents data for one syllable within a bird (n=13 syllables across the eight birds).

### Lack of relationship between experimental variation in motivational and performance aspects of song

The lack of difference between various aspects of LD and VD song performance contrasts with the difference in the motivation to produce LD and VD song. This suggests that the motivation to produce songs to live versus video presentations of females is independent of song performance. To further investigate the relationship between motivational and performance aspects of song, we assessed whether individual variation in the differential motivation to produce courtship songs to video versus live presentations of females correlated with individual variation in the differential modulation of performance features from UD (baseline) song to VD or LD song (first bouts only). Specifically, we calculated the difference in motivation as the difference in time total time spent singing LD and VD song and correlated this difference with the difference in performance modulation, measured as the difference in percent change from UD to LD (modulation when singing LD song) and from UD to VD song (modulation when singing VD song). Overall, we observed no significant correlations between experimental variation in motivation and performance (Figure 7; introductory notes: r=−0.15, p=0.7219; first motif duration: r=0.39, p=0.3380; FF of CV: r=0.51, p=0.1975).

**Figure 7:**
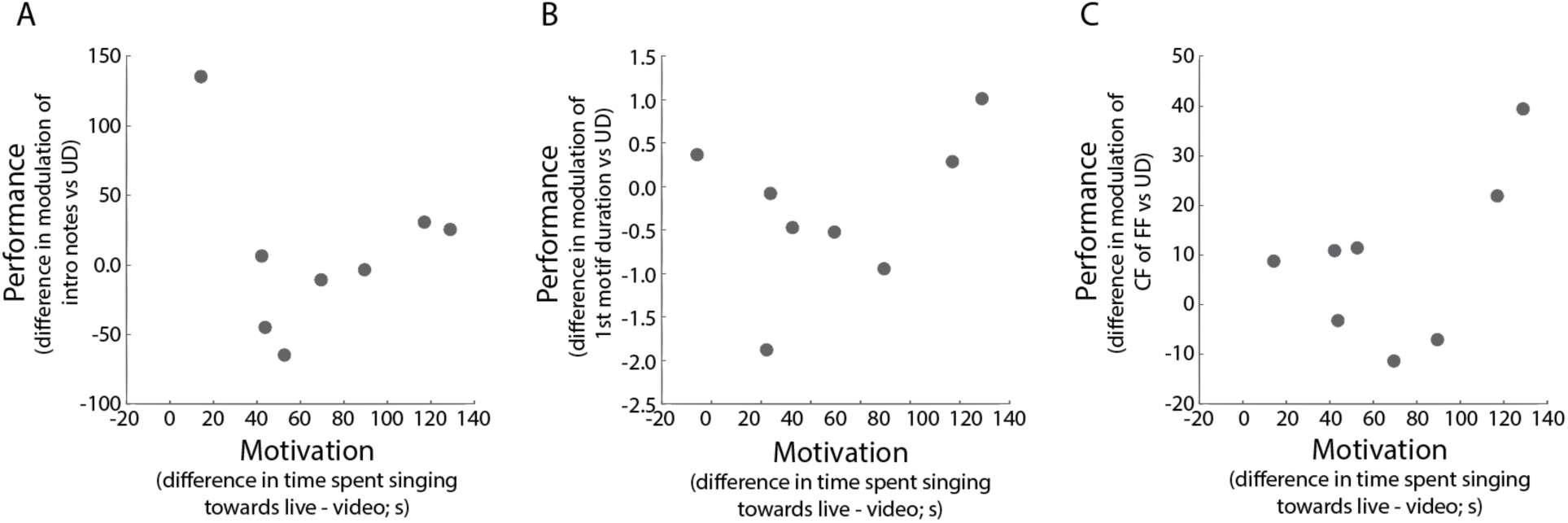
Relationships between measures of motivation and performance. For all panels, individual variation in motivation (difference in time spent singing LD and VD song; x-axis) is compared to individual variation in the modulation of song performance (differences in percent change of features from UD to VD or LD; y-axis). Because each bird only has one value for difference in motivation and because we measured the CV of FF for multiple syllables within a bird’s song, for this analysis we averaged the percent change in the CV of FF across syllables within each bird (i.e., one value per bird for CV of FF). Correlations were not significant for introductory notes (A), first motif durations (B) or the CV of FF (C).

## DISCUSSION

Male songbirds direct songs at females as part of their courtship ritual to secure copulations. This aspect of courtship can be analysed from both motivational and performance perspectives, with the former referring to the “drive” to produce courtship song and the latter referring to the acoustic features of courtship song (e.g., song tempo and stereotypy). Both aspects of courtship song are important because deficits in either component can affect attractiveness and mating success (Gil and Gahr, 2002; Heinig et al., 2014; Sakata and Vehrencamp, 2012; Woolley and Doupe, 2008). However, little is known about the extent to which these aspects of courtship song are regulated by similar or distinct mechanisms. Indeed, because neural circuits regulating the motivation to sing project to brain areas that regulate song performance (Riters, 2012), it is possible that motivational and performance aspects of courtship song are linked.

Here, we took advantage of previous studies that outline experimental manipulations that affect the motivation to produce courtship song and assessed the degree to which such motivational variation was associated with variation in vocal performance. Previous studies indicate that male songbirds will produce courtship songs to videos of females but tend to be less motivated to sing to videos of females than to live presentations of females (Galoch and Bischof, 2007; Ikebuchi and Okanoya, 1999; Takahasi et al., 2005). Consequently, we analysed whether the vocal performance of courtship songs produced to videos of females was distinct from songs produced to live females. Consistent with previous studies, we found that male zebra finches produced courtship songs to video presentations of female conspecifics but were less motivated to produce courtship songs to video presentations of females than live presentations of females. Specifically, males produced less the half the amount courtship song during video presentations of females than during live presentations, and this difference was due to males being less likely to produce courtship songs during video presentations and producing shorter songs when courting videos of females (Figure 3). However, courtship songs that male zebra finches produced to videos of females were structurally indistinguishable in most ways from courtship songs produced towards live females. In particular, the number of introductory notes preceding song, song tempo, and the variability of the fundamental frequency of syllables with flat, harmonic structure were not significantly different between video-directed (VD) and live-directed (LD) songs (Figure 5). Consequently, these data support the notion that the motivation to produce courtship song is controlled by independent mechanisms than the regulation of song performance, a notion that is further supported by the finding that individual variation in motivation to produce VD and LD songs was not related to individual variation in the modulation of VD and LD performance.

Such a dissociation between motivational and performance aspects of song has also been reported in previous studies (Cornil and Ball, 2010; Ritschard et al., 2011; Toccalino et al., 2016). For example, Toccalino et al. (2016) document that the familiarity of a female (i.e., repeated presentations of the same female) decreases the motivation of a male to direct courtship song to that female but does not affect performance aspects of his courtship song. In addition, Alward et al. (2013) found that testosterone implants into the medial preoptic area increased the amount of songs that male canaries produced to females but did not affect song performance measures such as song stereotypy. Collectively, these data suggest that distinct mechanisms contribute to motivational and performance aspects of birdsong and encourage experiments that further tease apart these aspects. Indeed, studies that revealed a dissociation between appetitive and consummatory aspects of copulatory behaviour (Moses et al., 1995; Pfaus et al., 1990; Riters et al., 1998; Seredynski et al., 2013) deeply shaped perspectives on social behavioural control and inspired a range of different experiments (Balthazart and Ball, 2007; Cornil et al., 2018).

Additionally, this interpretation suggests a need to revisit or build upon existing models of song motivation and control. Catecholamine (e.g., dopamine) release from midbrain and hindbrain circuits is hypothesized to contribute to the motivation to produce courtship song. For example, individual variation in the motivation to produce courtship song is correlated with variation in the number of dopamine-synthesizing neurons in the ventral tegmental area (VTA) of male zebra finches (Goodson et al., 2009), and manipulations of catecholaminergic neurons affect the likelihood that male zebra finches will produce courtship song to females (Barclay et al., 1996; Vahaba et al., 2013). Dopaminergic neurons in the VTA and periaqueductal grey (PAG) and noradrenergic neurons in the locus coeruleus (LC) project to various brain areas that regulate song control, include the avian basal ganglia nucleus Area X and the sensorimotor nucleus HVC (Appeltants et al., 2000; Castelino and Schmidt, 2010; Hamaguchi and Mooney, 2012; Maney, 2013; Tanaka et al., 2018), and dopamine or norepinephrine release into these areas affects neural activity and song performance (Cardin and Schmidt, 2004; Castelino and Ball, 2005; Ding and Perkel, 2002; Ihle et al., 2015; Leblois and Perkel, 2012; Leblois et al., 2010; Matheson and Sakata, 2015; Sasaki et al., 2006; Sizemore and Perkel, 2008; Solis and Perkel, 2006; Woolley, 2019). Taken together, this model suggests that variation in motivation should lead to variation in the amount of dopamine or norepinephrine released into areas like Area X or HVC, which should lead to variation in song performance. Our data do not support this model, suggesting that modifications or additional data are required. For example, further knowledge about the precise neural populations that regulate the repetition of introductory notes (e.g. Rajan and Doupe, 2013), song tempo (Long and Fee, 2008; Zhang et al., 2017) and the variability of fundamental frequency (reviewed in Woolley and Kao, 2015), and about the extent to which these populations receive catecholaminergic inputs would allow us to refine these models that link motivation and performance. Further, discovery of neurochemical systems that independently modulate song motivation or performance would greatly contribute to our understanding of this dissociation.

In addition to addressing models of vocal communication and social behaviour in songbirds, our results also extend previous studies in important ways by demonstrating that video presentations of female conspecifics lead to comparable changes to song performance as live presentations of females. The lack of significant differences in acoustic features between VD and LD song (Figure 5) and the correlations in the degree of vocal modulations when males directed songs at live or video presentations of females (Figure 6) indicate that videos of females are effective at eliciting the same suite of vocal performance changes as live presentations of females. From a mechanistic perspective, these data also suggest that videos of females engage the neural circuits for song performance to a comparable extent as live presentations of females. Neural activity in the anterior forebrain pathway (AFP) regulates context-dependent changes in the variability of fundamental frequency (reviewed in Brainard and Doupe, 2013; Murphy et al., 2017; Sakata and Vehrencamp, 2012; Woolley and Kao, 2015), whereas neural activity in the vocal motor pathway (VMP) has been proposed regulate context-dependent changes to temporal features of songs such as song tempo or the number of introductory notes before song (Aronov and Fee, 2012; Hampton et al., 2009; Matheson et al., 2016; Murphy et al., 2017; Rajan and Doupe, 2013; Stepanek and Doupe, 2010). Our data suggest that videos of females modulate neural activity in these circuits in the same way and to the same extent as live females.

The reason for differences in the motivation to produce courtship song to live vs. video presentations of females remains unknown. One possibility is that variation in female behaviour across conditions could account for this difference. For example, females in the videos were quiet and provided no real-time feedback to courting males. In contrast, although the behaviour of female stimulus animals was not quantified, live stimulus females can occasionally vocalize or posture during exposures to males. These behaviours could serve as feedback signals to the male and affect his motivation to produce song. As such, it is possible that male zebra finches perceived the females in the video as inattentive or uninterested in the male, which could have led to the male producing fewer and shorter songs towards these females (e.g. Ware et al., 2016). Given these differences, a useful next step would be to assess how female vocalizations and movements (e.g., using different types of videos of females) influence the motivation to produce courtship song in male zebra finches.

Broadly speaking, our results support the notion that video playbacks are a powerful tool to reveal the mechanisms by which individuals alter evolutionarily important behaviours, including vocal performance (Heinig et al., 2014; Podos et al., 2009; Sakata and Vehrencamp, 2012; Woolley et al., 2014). These findings also suggest that a standardized set of video stimuli can be used to reveal neural mechanisms underlying song motivation and performance (see Supplementary Information) and provide additional impetus to evaluate how specific visual and/or auditory information regulate song motivation and performance.

## ACKNOWLEDGEMENTS

We thank A. Jalayer for assistance with data analysis, and S.C. Woolley for constructive input and feedback throughout the experiment.

## COMPETING INTERESTS

No competing interests declared.

## FUNDING

This work was supported by funding from the National Science and Engineering Research Council (#05016 to J.T.S.); McGill University (Faculty of Science to R.F.), Canada Graduate Scholarship-Master’s (CGS-M to R.F.), Fonds de recherche du Québec – Nature et technologies, Master’s research scholarship (B1 to R.F.), and a Heller award (L.S.J.).

